# Pro-IL-18 secreted by keratinocytes detects the group A streptococcal protease SpeB

**DOI:** 10.1101/2021.08.22.457234

**Authors:** Anders F. Johnson, Jenna S. Sands, Keya Trivedi, Raedeen Russell, Doris L. LaRock, Christopher N. LaRock

**Affiliations:** Department of Microbiology and Immunology and Department of Medicine, Emory School of Medicine, Atlanta, GA, United States of America

## Abstract

Group A *Streptococcus* (GAS, *Streptococcus pyogenes*) is a professional human pathogen that commonly infects the skin. Keratinocytes are one of the first cells to contact GAS, and by inducing inflammation, they can initiate the earliest immune responses to pathogen invasion. Here, we characterized the proinflammatory cytokine repertoire produced by primary human keratinocytes and surrogate cell lines commonly used *in vitro*. Infection induces several cytokines and chemokines, but keratinocytes constitutively secrete IL-18 in a form that is inert (pro-IL-18) and lacks proinflammatory activity. Canonically, IL-18 activation and secretion are coupled through a single proteolytic event that is regulated intracellularly by the inflammasome protease caspase-1 in myeloid cells. The pool of extracellular pro-IL-18 generated by keratinocytes is poised to sense extracellular proteases and is directly processed into a mature active form by SpeB, a secreted GAS protease that is a critical virulent factor during skin infection. This mechanism contributes to the proinflammatory response against GAS, resulting in T cell activation and the secretion of IFN-γ that restrict GAS growth. Other major bacterial pathogens and microbiota of this skin did not have significant IL-18-maturing ability. Taken together, these results suggest keratinocyte-secreted IL-18 is a sentinel that sounds an early alarm that is highly sensitive to GAS, yet tolerant to non-invasive members of the microbiota.

## Introduction

The skin provides the first resistance to infectious, chemical, and physical insults. The obligate human pathogen Group A *Streptococcus* (GAS, *Streptococcus pyogenes*) colonizes oropharyngeal mucosa and epidermal surfaces, specifically adhere to and invading skin epithelial cells and keratinocytes [1–4]. Beyond superficial infections like impetigo, further tissue invasion can lead to cellulitis, sepsis, and necrotizing fasciitis. Invasive infections, and immune-mediated sequalae like rheumatic heart disease, are responsible for an estimated half a million annual deaths globally [5]. These infections are highly inflammatory, which contributes to their pathology and complicates their treatment, and several lines of evidence suggest GAS use this inflammation to promote replication and transmission [6–9]

Interleukin-18 (IL-18) is a proinflammatory cytokine that induces cell-mediated immunity [10]. IL-18 is detected by the IL-18R/IL-18RAP (IL18R1/IL1R7, IL-18Rα/IL-18Rβ) receptor complex and works with IL-12 to induce IFN-γ and Th1-type responses from T cells, NK cells, and dendritic cells [11–13]. Accordingly, IL-18 is important in the host defense against *Salmonella* Typhimurium [14], *Shigella flexneri [15], Yersinia enterocolitica* [16], *Listeria monocytogenes* [17], *Burkholderia pseudomallei* [18], *Mycobacterium tuberculosis* [19], *Streptococcus pneumoniae* [20], and group B *Streptococcus* [21]. Newly-synthesized IL-18 (pro-IL-18) is inert and requires removal of an amino-terminal autoinhibitory domain, canonically by the host protease caspase-1 [11,12]. Caspase-1 is itself regulated by the inflammasome complex, which in myeloid cells can also regulate IL-1β maturation and the cell death program pyroptosis [22]. Caspase-8 [23], granzyme B [24], chymase [25], proteinase 3 [26], and neutrophil elastase [27] also cleave pro-IL-18, though their physiologic relevance in IL-18 activation remains to be established. Human keratinocytes constitutively express and release pro-IL-18 [28–31], suggesting the skin as an anatomical location where these or other extracellular proteases may participate in IL-18 activation.

We hypothesized that GAS interactions with human keratinocytes contributes to the proinflammatory storm observed during infection. This study shows that keratinocytes release numerous proinflammatory cytokines during GAS infection, including pro-IL-18, which is directly matured by the SpeB cysteine protease of GAS. This mechanism supports a model wherein SpeB acting as a “bacterial caspase” that proteolytically activates proinflammatory cytokines of the IL-1 family, which in term may limit GAS invasive infection.

## Results

### GAS induces keratinocytes secretion of proinflammatory cytokines

The epithelium is one of the first body tissues GAS will contact. Therefore, the resident cells provide one of the earliest responses to infection by producing antimicrobial effectors and proinflammatory cytokines and chemokines that coordinate immune response [32–35]. To assess the cytokine repertoire elicited by GAS, we infected cultured and primary cells epithelial cells with M1T1 GAS 5448, a highly virulent strain associated with modern epidemic invasive infections [36]. Similar to previous observations [32,33,37], we observed robust secretion of MIF, IL-8, and other proinflammatory cytokines and chemokines by infected HaCaT keratinocytes (**Figure 1a**). Detroit 562 human pharyngeal epithelial cells, HEp-2 human laryngeal epithelial cells, and A-431 human keratinocytes all secreted a similar, nonredundant, cytokine profile. However, compared to each of these cell lines, human primary keratinocytes expressed a greater diversity in CXC-family chemokines that were absent with any individual cell line and the pro-inflammatory cytokine IL-1α (**Figure 1a**). In contrast, human primary endothelial cells also secreted cytokines and chemokines but had a distinct repertoire. No cell produced detectable CCL1, CCL5, G-CSF, IFN-γ, IL-1β, IL-2, IL-4, IL-5, IL-10, IL-12, IL-13, IL-16, IL-17A, IL-17E, IL-21, IL-27, IL-32α, MIP-1,TNF-α. Primary keratinocytes and endothelial cells each formed a distinct group relative to all cell lines by multivariate analysis (**Figure 1b**), due to each secreting a distinct cytokine profile (**Figure 1c**).

**Figure 1.**
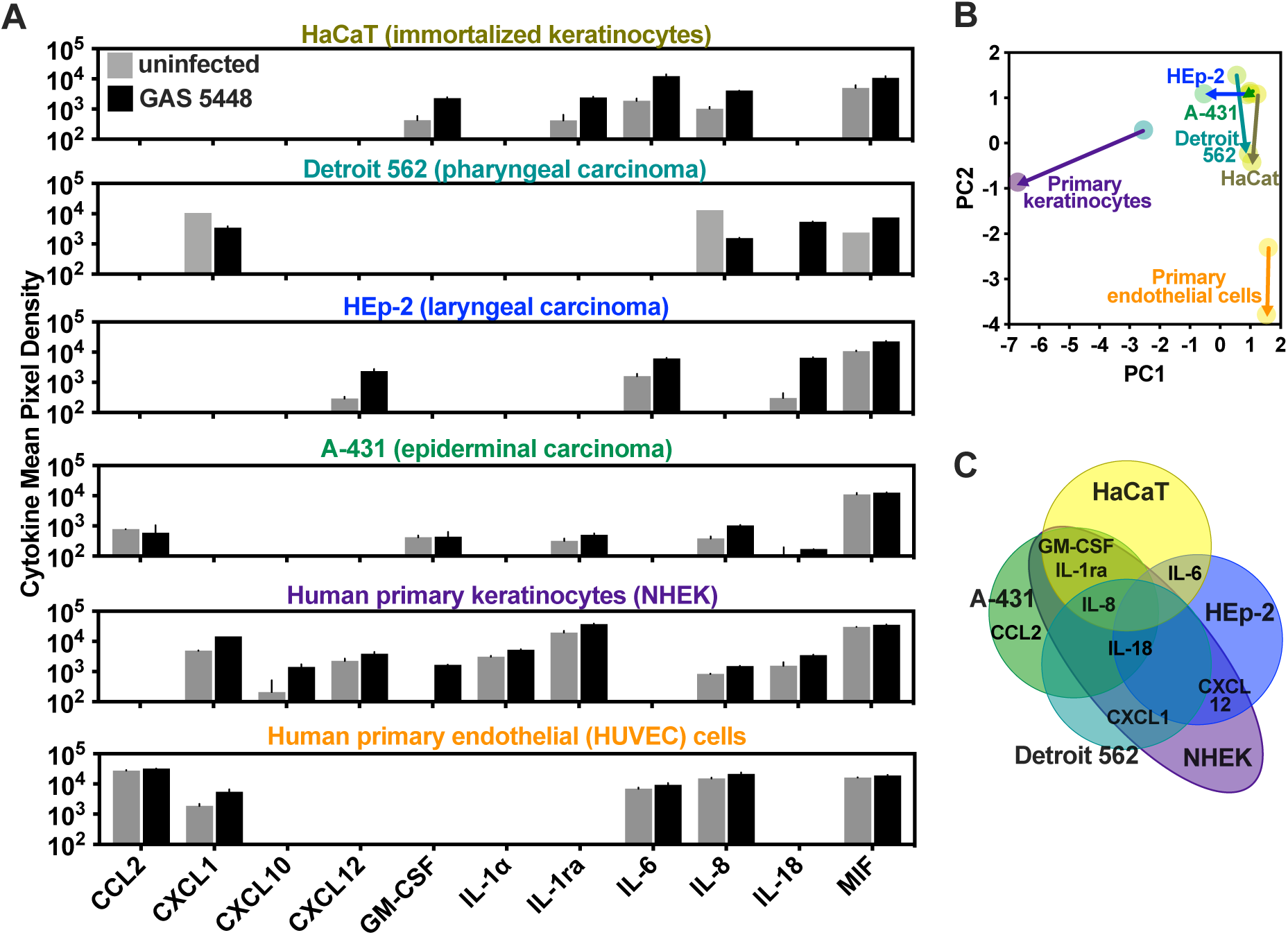
Cytokine profiles of keratinocytes and related cell lines. HaCaT, Detroit 562, HEp-2, A-431, primary keratinocytes, or HUVEC cells were infected with 7.5 × 10^6^ colony forming units (CFU) of GAS for 6 h. (**A**) Relative abundance of select cytokines was examined by membrane-based antibody array. (**B**) Cytokine profiles of each cell were examined by multivariate (principal component analysis). Arrows indicate change in cells from uninfected to 6 h infection. (**C**) Graphical representation of the congruent cytokine profiles between cell types.

### GAS activates IL-18

One cytokine of particular interest to us was IL-18, an IL-1 family cytokine that bridges innate and adaptive immune responses stimulate IFN-γ production by T and NK cells [10]. In immune cells, IL-18 release is conventionally regulated by the inflammasome protease caspase-1 alongside the cell death program, pyroptosis [22]. In contrast, healthy keratinocytes have been seen to constitutively release IL-18 [29], consistent with our observation that GAS infection was not required for IL-18 secretion (**Figure 2a**). Since chronic IL-18 signaling would be pathological, we next examined whether the activity of IL-18 secreted by keratinocytes. The determinant of activity for cytokines that are post-translationally regulated by proteolysis is not necessarily specific to a single regulator, and is exclusively whether the product formed is able to bind its cognate receptor and stimulate receptor signaling [6]. To measure whether keratinocyte-released IL-18 had biological activity, we used HEK-Blue™ IL-18 Reporter Cells, which have been engineered to express the IL-18 receptor complex (IL-18R and IL-18RAP) and induce an alkaline phosphatase reporter in response to IL-18, but not other cytokines or pathogen-associated molecular patterns. None of the IL-18 released by uninfected keratinocytes had activity, however, it was rendered active during GAS infection (**Figure 2b**). Next, we took cell-free supernatants from keratinocytes that contained this immature form of IL-18 and incubated it with GAS. GAS was able to again convert IL-18 into an active form (**Figure 2c**). Since IL-18 activation occurred in the absence of keratinocytes, we concluded that conventional cellular regulators (the inflammasome) were not essential, and instead, one or more GAS factors directly act on inert pro-IL-18 in a manner that renders it active.

**Figure 2.**
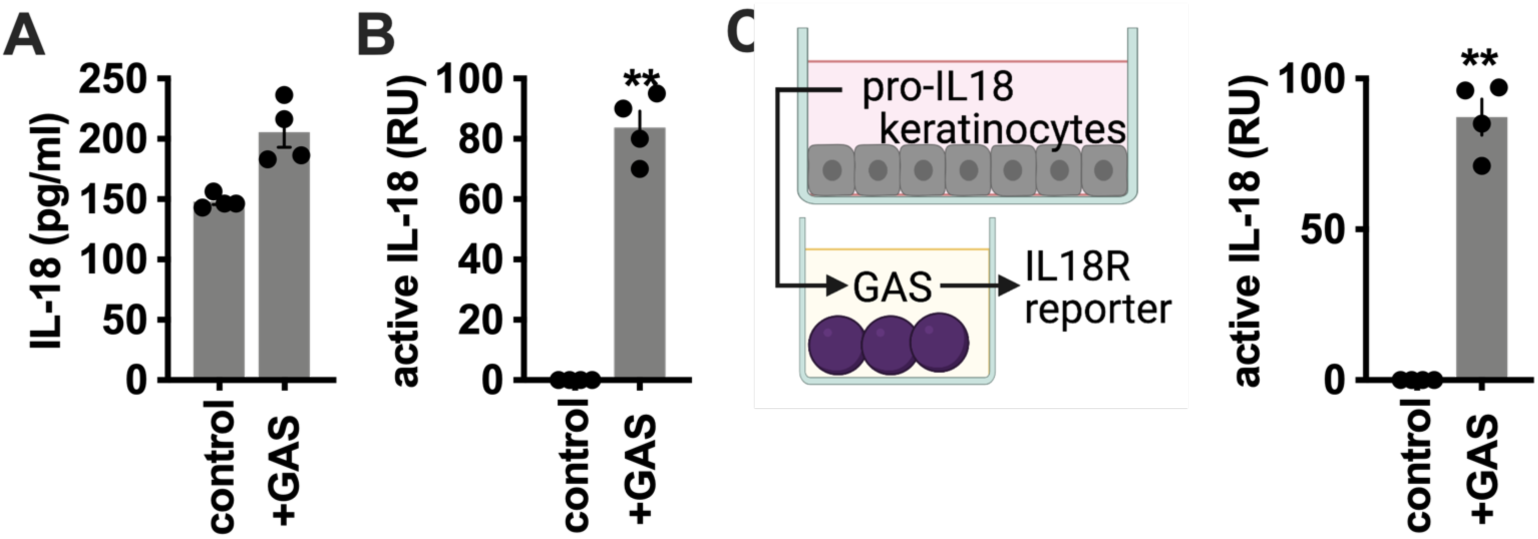
Examination of IL-18 activation in GAS-infected keratinocytes. Primary human keratinocytes were infected with GAS at a multiplicity of infection (MOI) of 10 for 4 h. (**A**) Secreted IL-18 (total, pro-and mature forms) was measured by ELISA. (**B**) Bioactive IL-18, capable of inducing signaling through the IL-18R/IL-18RAP receptor complex, was measured with HEK-Blue IL18 reporter cells. (**C**) Supernatants removed from primary human keratinocytes were incubated with GAS and IL-18 activity measured as in (B). Data were analyzed by 1-way ANOVA using Dunnett multiple comparisons analysis. All data represent at least 3 independent experiments with 4 replicates. Bars show median values ± standard deviation. \*\**P* < .005; ns, not significant.

### The GAS protease SpeB activates pro-IL-18

In order to determine the bacterial determinants required for direct conversion of pro-IL-18 by GAS, we screened a panel of defined virulence factors mutants. During keratinocyte infection, we found that expression of the secreted cysteine protease SpeB was essential for GAS to convert IL-18 to an active form (**Figure 3a**). Consistent with this observation, CovRS, a two-component regulator of SpeB that is frequently inactivated in pathoadapted GAS that evade immune restriction, was also required for GAS conversion of IL-18 (**Figure 3b**). Plasmid complementation of Δ*speB* GAS 5448 restored activation, but plasmid encoding the catalytically inactive form SpeBC192S did not, indicating the protease activity of SpeB was required for IL-18 conversion (**Figure 3c**). We tested gain-of-function using *Lactococcus lactis* made to express SpeB and found that the catalytically active SpeB was necessary and sufficient for IL-18 activation (**Figure 3d**). Furthermore, in the absence of infection or other treatment, purified SpeB was sufficient to activate IL-18 (**Figure 3e**).

**Figure 3.**
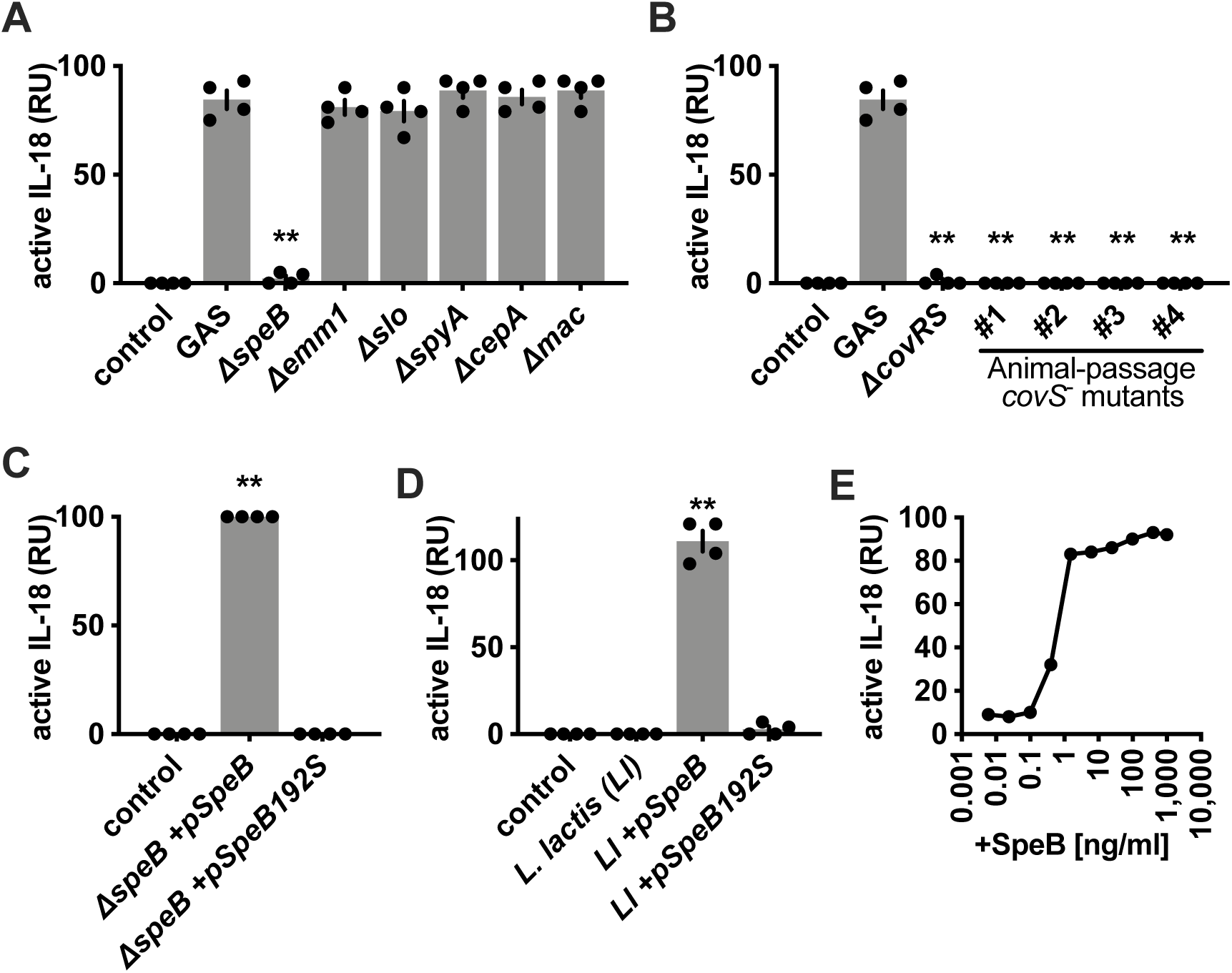
Examination of GAS requirements for IL-18 activation. Primary human keratinocytes were infected with GAS at MOI 10 for 4 h, then bioactive IL-18 was measured with HEK-Blue IL-18 reporter cells. (**A** and **B**) IL-18 activation was measured in the supernatants from keratinocytes infected with the indicated gene knockouts of GAS strain 5448. (**C**) IL-18 activation was measured in the supernatants from keratinocytes incubated with spectinomycin and anhydrotetracycline to maintain SpeB expression during infection with Δ*speB* GAS 5448 carrying the indicated plasmids. (**D**) IL-18 activation was measured in the supernatants from keratinocytes during infection with *L. lactis*, spectinomycin and anhydrotetracycline were included during infections with the plasmid-carrying strains. (**E**). IL-18 activation was measured in the supernatants from keratinocytes treated with titrations of purified SpeB protein. Data were analyzed by 1-way ANOVA using Dunnett multiple comparisons analysis. All data represent at least 3 independent experiments with 4 replicates. Bars show median values ± standard deviation. \*\**P* < .005; ns, not significant.

### pro-IL-18 is matured by SpeB cleavage

IL-18 is conventionally activated when an N-terminal pro-domain (pro-IL-18) is proteolytically removed by Caspase-1 [22]. The truncated C-terminal portion then is freed from the steric hindrance that blocks binding and formation of a signaling complex with IL-18R and IL-18RAP [38]. In contrast to Caspase-1, which converted the ∼24 kDa pro-IL-18 precursors to a ∼18 kDa product, SpeB generated larger products spanning ∼22 to 20 kDa in size (**Figure 4a**). This is similar to IL-1β, which SpeB cleaves at a different site than Caspase-1 to form a larger product that nonetheless is active [6]. However, there is no significant homology between the prodomains of IL-18 and IL-1β in mice or humans. The region proximal to the Caspase-1 cleavage site (after amino acid 36) of IL-18 makes contacts with the IL-18R in the co-crystal structure that stabilize binding and likely contribute to avidity and signaling [39]. Since the N-terminus occludes these interactions when attached [38], we were intrigued by how N-termini of different lengths generated by different protease cleavage sites impact IL-18 signaling (**Figure 4b**). Thus, we next generated a series of N-terminal truncations by *in vitro* transcription-translation to examine their activity with IL-18 reporter cells. Unlike IL-1β, which similar techniques showed is only activated within a discrete internal region [40], truncation of as few as ten amino acids from the N-terminus of IL-18 was sufficient to significantly increase signaling (**Figure 4c**). Products smaller in size than those generated by Caspase-1 rapidly loose activity, suggesting these residues are important in receptor interaction, but larger products like those generated by SpeB retained significant signaling activity when generated in recombinant forms (**Figure 4c**).

**Figure 4.**
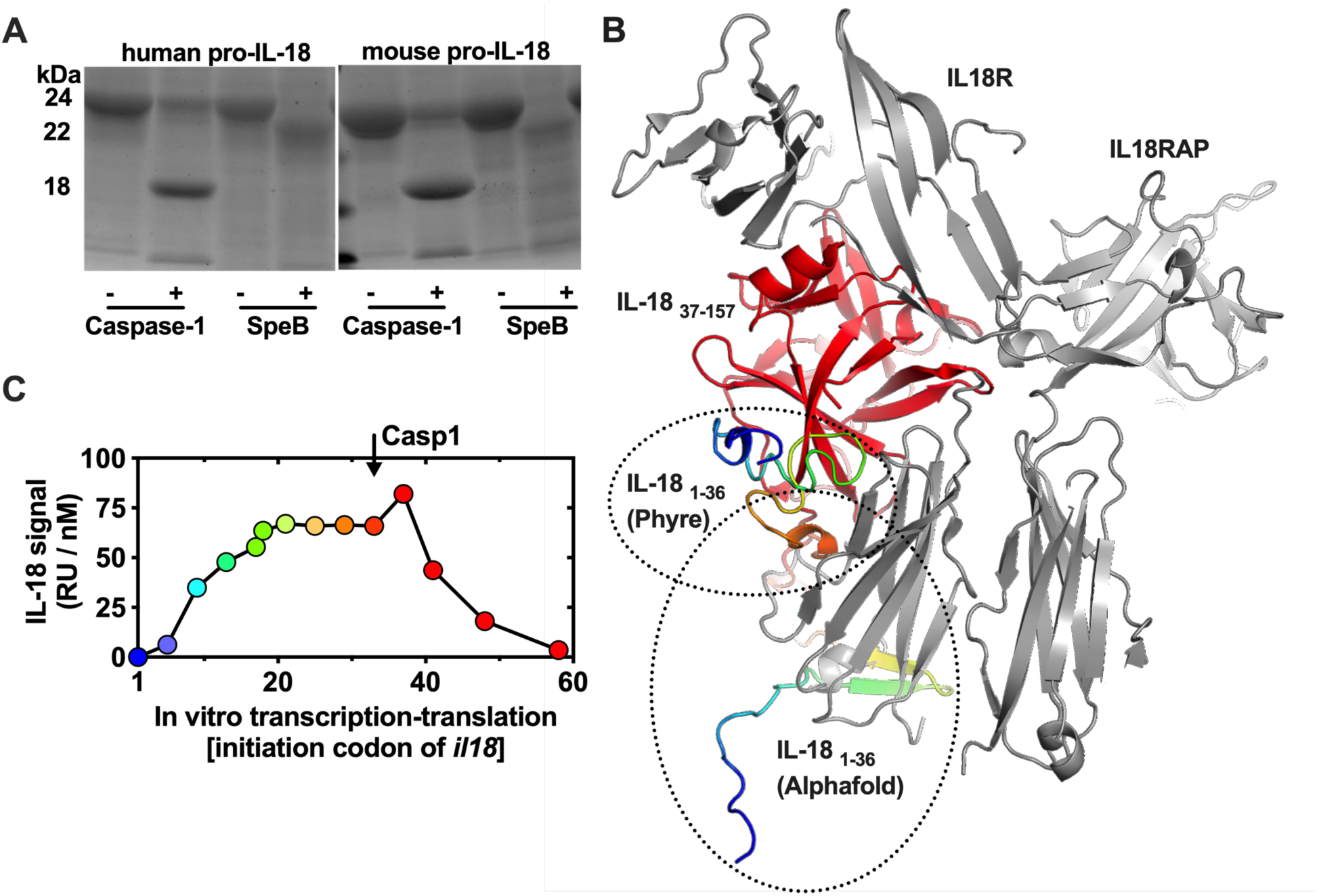
Examination of SpeB processing of IL-18. (**A**) Recombinant human and murine pro-IL-18 was purified and incubated with purified, active human Caspase-1 or purified, active SpeB, then cleavage products separated by SDS-PAGE and visualized by staining. (**B**) Structure of the human IL-18 receptor complex (PDB:3wo4; [39]), IL18R and IL18RAP; grey, residues 37-157 of IL18 (caspase-cleaved product); red, predicted N-terminal (residues 1-36) inhibitor domain; rainbow. (**C**) Signaling activity of recombinant IL-18 N-terminal truncations generated using *in vitro* transcription-translation from the human *il18* gene with coding beginning at the indicated codon, 1 is full-length pro-IL-18; colors match positions in (B). Activity toward HEK-Blue IL18 reporter cells was normalized for loading by quantifying total product using IL-18-specific ELISA.

### SpeB-activated IL-18 induces inflammation

IL-18 was originally discovered as a factor that induces type II interferon (IFN-γ) production from Th1 cells [41]. Since IL-18 is elevated in experimental murine infections as it is during human infection [6,42,43], we decided to examine the role of IL-18 in a murine skin infection model. No difference in GAS growth was observed between wild-type C57Bl/6 and *il18*^-/-^ mice (**Figure 5a**). Thus, irrespective of whether caspases, SpeB, or other mechanism activate IL-18, this model would be unable to capture whether that specific mechanism would have protective benefit. Nevertheless, IL-18 are IFN-γ and critical in human immunity and are protective against infection [44–46]. Only a few instances have been identified where *il18*^-/-^ mice are susceptible to infection [19,47–49], suggesting differences in immune signaling between species or methods for modeling infection fail to capture IL-18-mediated signaling. IL-18 and T cell responses are instead commonly approached using human primary cells such whole blood or PBMCs to get around limitations imposed by mice [43,50–53]. Inhibition of IL-18 or IFN-γ with neutralizing antibodies promoted GAS growth in whole human blood (**Figure 5b**). While this does not specifically show a role for keratinocyte-secreted IL-18 in the immune response, it supports that these cytokines can have a protective role of in the response of human immune cells to GAS, as they do for other pathogens.

**Figure 5.**
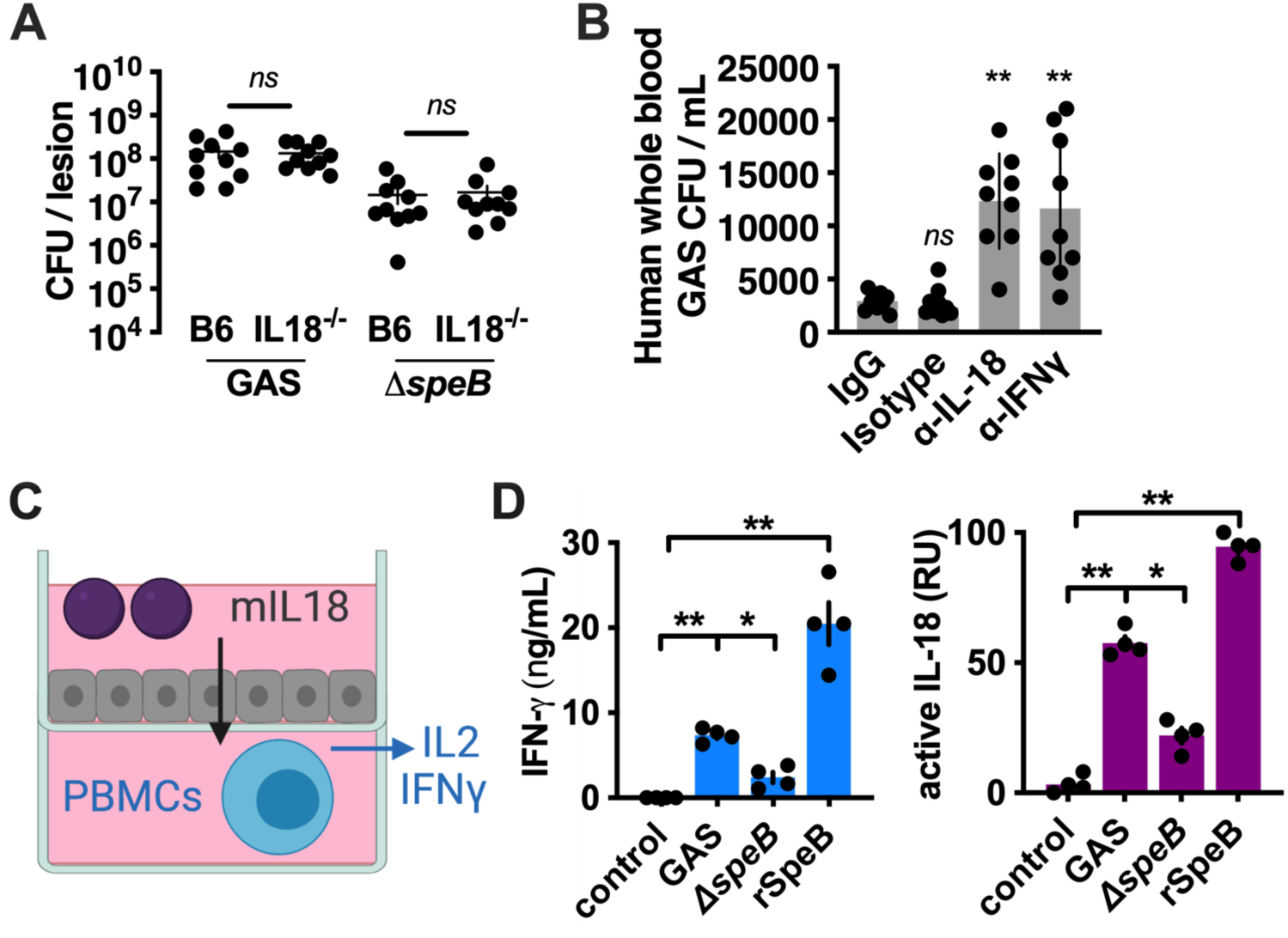
SpeB activation of IL-18 promotes antimicrobial IFN-γ responses. (**A**) C57BL/6 wild-type or IL18-knockout (IL18^-/-^) mice were inoculated intradermally with 10^8^ CFU of GAS 5448 or its Δ*speB* mutant. After 72 h, mice were euthanized and GAS CFU enumerated at the infection site. Results are from 2 independent experiments with 5 mice in each. (**B**) Heparinized whole human blood treated with isotype antibody control, α-IL-18, or α-IFN-γ neutralizing antibodies (50 µg/ml) was inoculated with 10^5^ CFU of GAS 5448 and CFU enumerated after 4 h. Results are pooled from 3 independent experiments. (**C**) Diagram of primary keratinocyte/PBMC co-culture model; IFN-γ is a reporter of T cell activation, which can occur via IL-18 and additional mechanisms during infection. (**D**) Cocultured keratinocyte/PBMCs were infected with 10^5^ CFU of GAS or treated with rSpeB. After 4 h, active IL-18 was quantified with HEK-Blue IL-18 reporter cells, and IFN-γ by ELISA. Data represent at least 3 independent experiments with 4 replicates. Data were analyzed by 1-way ANOVA using Dunnett multiple comparisons analysis. Bars show median values ± standard deviation. \**P* < .05; \*\**P* < .005; ns, not significant.

PBMCs produce IFN-γ in response to mature IL-18 when cultured with IL-12 [41]. Thus, we further examined whether IL-18 released by keratinocytes and processed by GAS is sufficient to induce IFN-γ release in a co-culture model where keratinocytes are incubated with PBMCs and then infected with GAS (**Figure 5c**). SpeB-expressing GAS induced significantly greater IFN-γ, and purified SpeB was sufficient to induce IFN-γ (**Figure 5d**). IFN-γ production correlated with the production of active IL-18 (**Figure 5d**). IL-18 was no longer fully SpeB-dependent as in keratinocyte monoculture (**Figure 2b**), but was nonetheless significantly enhanced by SpeB (**Figure 5d**).

### IL-18 activation is species-restricted

Proteases are one of the most common protein classes and are frequently employed by pathogens as virulence factors. Like IL-18, IL-1β is directly cleaved and activated by bacterial proteases other than SpeB, such as *Pseudomonas aeruginosa* LasB [6,40]. These proteases potentially represent a group of “bacterial caspases” that may share in common the activation of these two related IL-1 family cytokines. We adapted our earlier models of keratinocyte infection (**Figure 2b**) and cell-free processing of secreted pro-IL-18 (**Figure 2c**) to screened for species that can directly activate IL-18 (**Figure 6a**). We focused on a panel of Streptococci, many commonly reside on the human epithelia of the skin and upper respiratory tract, either as microbiota or pathogens, where they may contact keratinocyte-secreted pro-IL-18. During co-culture where live bacteria could infect live keratinocytes, nearly all species induced IL-18 activation by keratinocytes, which could proceed via conventional host processes, ie. the inflammasome (**Figure 6b**). However, only GAS and *S. mutans* significantly activated the inert pool of pro-IL-18 in cell-free supernatants collected from naïve keratinocytes (**Figure 6b)**. This suggests constitutive pro-IL-18 secretion does not result in chronic inflammation in part because few proteases convert convert IL-18 into an active form.

**Figure 6.**
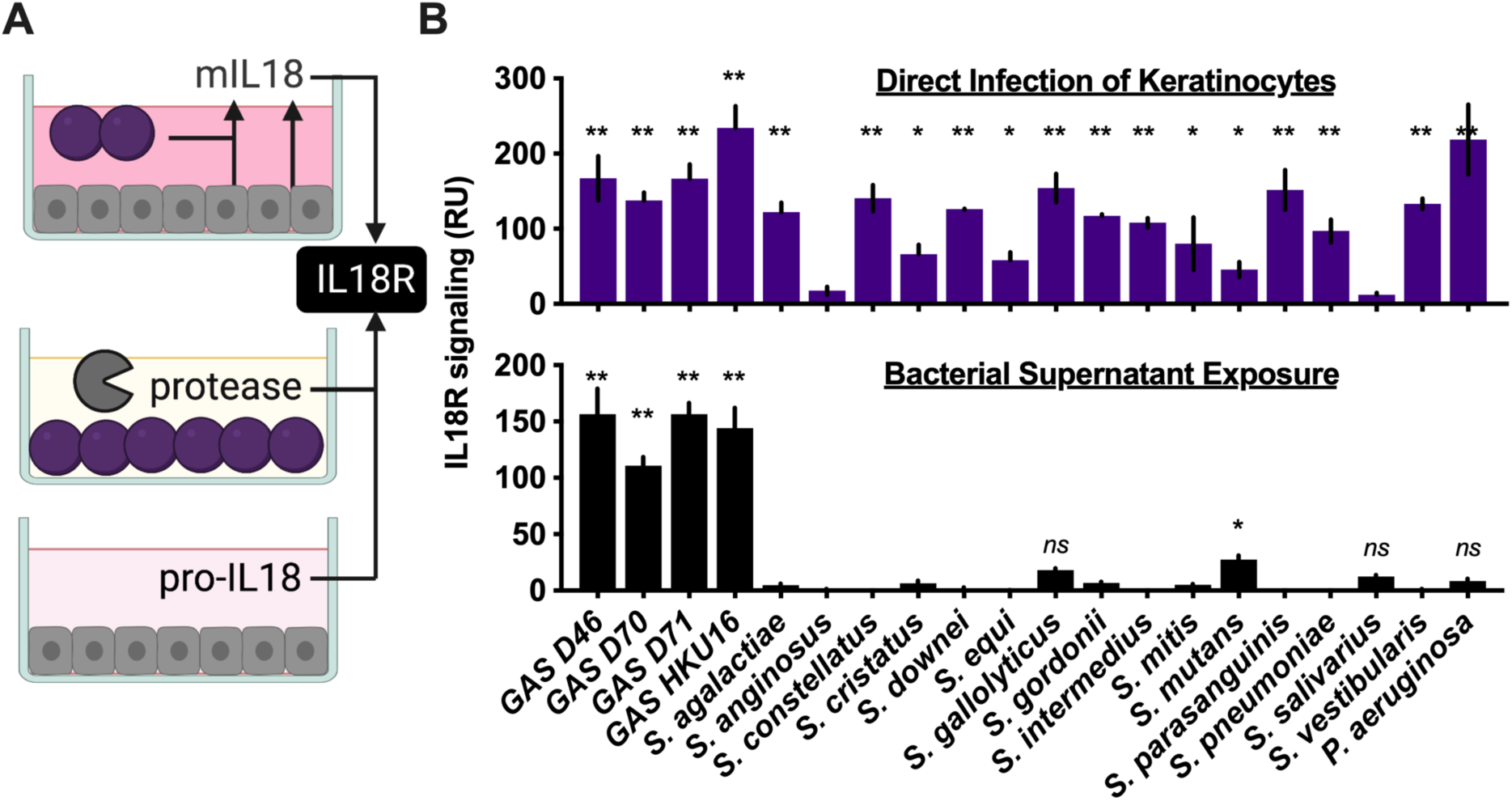
Bacterial activation of IL18. (**A**) Diagram of infection model. (**B**) In ‘Direct Infection of Keratinocytes’ primary human keratinocytes were infected with each bacteria at MOI 10. In ‘Bacterial Supernatant Exposure’, an equivalent volume of overnight culture supernatant was combined with cell-free supernatants from keratinocyte *ex vivo*. After 4 h, for both models, bioactive IL-18 was measured with HEK-Blue IL18 reporter cells. Data represent at least 3 independent experiments with 4 replicates. Data were analyzed by 1-way ANOVA using Dunnett multiple comparisons analysis compared to uninfected/untreated keratinocytes. Bars show median values ± standard deviation. \**P* < .05; \*\**P* < .005; ns, not significant.

## Discussion

SpeB is important for GAS colonization throughout the body and for penetrating deeper into tissue during invasive infection [6,54]. Skin keratinocytes are thus poised to mount some of the earliest immune responses against this important pathogen, but must safeguard against aberrant activation by the microbiota, since they are in constant contact with numerous other species they must tolerate. Consistent with prior observations [28–31], we observed constitutive pro-IL-18 secretion by primary human keratinocytes (**Figure 1a, 2a, 2b**), which is directly activated by SpeB (**Figure 3e**) to induce proinflammatory and antimicrobial responses (**Figure 1a, 2a, 2b**). Members of the healthy skin microbiota do not similarly activate immature pro-IL-18 (**Figure 6b**), so we hypothesize this pool of cytokine poised for activation serves as an early sentinel for infection by potentially invasive pathogens that make use of proteolytic virulence factors. By directly sensing specific proteases required for infection, IL-18 allows keratinocytes to discriminate between numerous species with high virulence potential and common pathogen-associated molecular patterns and toxins. While GAS is readily detected, low-level activation by the cariogenic opportunistic pathogen *S. mutans* suggests this mechanism could have importance in dental health. Furthermore, there may be pathogens from genera other than the *Streptococcus* with this activity, since proteases are broadly used as virulence factors by many human pathogens.

The mechanism underlying IL-18 discrimination of proteases remains unclear. Recent NMR studies show that mature (caspase-1-cleaved) IL-18 has a distinct spectra from pro-IL-18, suggesting that receptor binding is inhibited by intramolecular interactions from the pro-domain inducing major structural changes in the cytokine [38]. Amino-terminal truncations showed removal of the first 8 amino acids gave folding similar to pro-IL-18, the first 10 an intermediate form, and 12, 13, or 22 a form resembling the mature protein [38]. This closely coincides with our observations, where our mapping by IVTT observes partial activity starting with a 9 amino acid truncation (**Figure 4c**). Alternative endogenous proteases results in a variety of sizes [24–27]. Our observations, with those of Tsutsumi et al. [38], suggest that proteases that remove even a small portion of the inhibitory N-terminus result in an active cytokine, whereas ones that generate smaller IL-18 forms may instead inactivate the cytokine. Nonetheless, the relatively loose substrate specificity of *Pseudomonas* LasB [40], with several potential cleavage sites in the N-terminus, did not result in cell-independent IL-18 activation (**Figure 6b**). Therefore, additional factors may regulate IL-18 activation beyond the primary sequence in this region, such as stabilizing reactions in the secondary structure burying side chains recognized by LasB or other promiscuous proteases in the environment.

We further observed primary keratinocytes secrete a more diverse set of cytokines relative to immortalized cell lines, including IL-18, a cytokine that serves as a crucial bridge from the innate immune responses to the adaptive immune cells. However, all cytokine profiles were variable between cell lines, which are commonly used or interpreted interchangeably. Autocrine or paracrine effects of these cytokines should be considered when there is potential to impact experimental findings. Furthermore, antimicrobial effectors are commonly co-regulated by many of the same signaling transduction networks, and these potential defects in each line can potentially mask pathogen phenotypes. Altogether, we reiterate caution should be exercised when immortalized cells are used for modeling host-pathogen interactions. Unlike keratinocytes, endothelial cells did not release IL-18. Therefore, extracellular activation of IL-18 by SpeB may not necessarily occur during sepsis or at distal sites from the skin surface, and may be most important in priming responses early in infection.

## Materials and Methods

### Bacterial Strains

GAS strain 5448, representative of the pandemic M1T1 clone, and its isogenic *ΔspeB, Δemm1, Δslo, ΔspyA, ΔcepA, Δmac, ΔcovRS*, and complemented pSpeB strains have been previously described [6,35,55]. GAS was statically grown at 37°C in Todd Hewitt broth (THB, Difco), washed two times with phosphate-buffered saline (PBS), and diluted to a multiplicity of infection (MOI) of 10 or 100 for *in vitro* infections. Selection for pSpeB was maintained with spectinomycin (Sigma) 100 ug/ml and expression controlled with titrations of anhydrotetracycline (Cayman Chemical) as previously [55]. The following reagent was obtained through BEI Resources, NIAID, NIH: *Streptococcus anginosus* F0211, *Streptococcus cristatus* F0329, *Streptococcus downei* F0415, *Streptococcus equi* ATCC 9528, *Streptococcus gallolyticus* TX20005, *Streptococcus intermedius* F0413, *Streptococcus mitis* F0392, *Streptococcus parasanguinis* F0449, and *Streptococcus vestibularis* F0396. *Streptococcus mutans* JH1140 was provided by S. McBride and *Pseudomonas aeruginosa* PA01 was used as previously described [40]. All additional strains are clinical isolates provided through the Emory University Investigational Clinical Microbiology Core. All bacteria were routinely propagated statically at 37°C in Todd Hewitt broth (Difco), washed two times with phosphate-buffered saline (PBS), and diluted to a multiplicity of infection (MOI) of 10 or 100 for in vitro infections.

### Cell Culture

Primary human keratinocytes were supplied by PromoCell and cultured in Growth Medium 2 with supplement (C-20011, C-39011; PromoCell). Primary umbilical vein endothelial cells were supplied by Lifeline Cell Technology and cultured in endothelial cell growth medium (C-22010; PromoCell). A-431 lung epidermal cells (ATCC), Detroit 562 pharyngeal cells (ATCC), HaCat cells (unavailable from a repository, provided by C. Quave, Emory), and HEp-2 laryngeal cells (ATCC) were cultured in Eagle’s Minimum Essential Medium (Gibco) supplemented with 10% heat-inactivated fetal bovine serum (hiFBS, Atlanta Biologicals). HEK-Blue™ IL-18 Reporter cells (Invivogen) were cultured in Dulbecco’s Modified Eagle’s Medium (Gibco) with 10% hiFBS. 100 U/mL penicillin, 100 µg/mL streptomycin, and 100 µg/mL Normocin was supplemented during routine culture and omitted during experimental infections, unless otherwise noted. All cells were maintained at 37 °C and 5% CO_2_.

### Cytokine Measurements

The relative levels of selected human cytokines and chemokines was determined in parallel using a membrane-based antibody array (ARY005B, R&D Systems). Cells were seeded at 7.5 × 10^5^ cells per well in a tissue-culture treated six-well plate. Cells were infected at an MOI of 10, and bacteria spun onto cells at 160 × g for three minutes. 1 mL of cell supernatant was collected after six hours of infection and incubated on the membrane overnight at 4°C and developed following the manufacturer’s protocol. Chemiluminescence was measured using the ChemiDoc MP imaging system, and mean pixel density was quantified using ImageJ. Only proinflammatory cytokines made by at least one of the cells lines under study are presented graphically. IL-18 signaling activity was measured in 50 µL volumes of cell supernatant using HEK-Blue™ IL-18 Reporter cells (Invivogen) and secreted alkaline phosphatase activity measured after 18 h incubation as previously [56]. This was normalized total IL-18 measured by ELISA (R&D Systems).

### Protein Purification

The fully spliced coding sequence for human *il18* and murine *il18*, encoding pro-IL-18 for each species, was generated by gene synthesis for expression from pETPP [35] to generate pETPP-hIL18 and pETPP-mIL18. Expression was induced from the plasmids in BL21 cell induced with 0.2 mM IPTG (Sigma) overnight at 18°C. Bacterial cell pellets were suspended in 10 mL of phosphate buffered saline (PBS, pH 7.4). Cells were completely lysed by sonication at 40% amplitude for two minutes for 30 seconds at 10 second intervals and centrifugation at 7500 × g for 10 minutes. Lysate was through Talon gravity columns loaded with HisPur™ Cobalt Resin (Thermo Scientific), washed with PBS, and eluted in PBS supplemented with 300 mM imidazole. SpeB was purified as previously [6]. Cleavage of purified human and murine pro-IL-18 was performed as previously with pro-IL-1β [6], with SpeB (100 ng) and Caspase-1 (1U, Novus) in PBS with 2 mM dithiothreitol 2 h at 37 °C. Reactions were stopped by the addition Laemmli buffer, 10% β-ME, and 1 × SDS loading buffer solution, then boiled at 95 °C for 5 min. Samples were analyzed by SDS-PAGE on Tris-Glycine gels (Invitrogen) and visualized with AquaStain (Bulldog Bio).

### In vitro transcription/translation

IL-18 truncations were generated through *in vitro* transcription/translation using pET-pro-IL-18 as a template in 10 µl reaction volumes following the manufacturer’s recommendations (TNT Coupled Reticulocyte Lysate; Promega) as previously [40], with 5’ primers gaaattaatacgactcactatagggagaccccacc-atggctatggctgctgaaccagtaga (IVTTIL18-1), gaaattaatacgactcactatagggagaccccaccatggctccag-tagaagacaattgcatc (IVTTIL18-5), gaaattaatacgactcactatagggagaccccaccatggctaattgcatcaactttgtggc (IVTTIL18-9), gaaattaatacgactcactatagggagaccccaccatggcttttgtggcaatgaaattta (IVTTIL18-13), gaaattaatac-gactcactatagggagaccccaccatggcttttattgacaatacgcttta (IVTTIL18-18), gaaattaatacgactcactatagggagaccccac-catggctaaatttattgacaatacgct (IVTTIL18-17), gaaattaatacgactcactatagggagaccccaccatggctaatacgctttactttatagc (IVTTIL18-21), gaaattaatacgactcactatagggagaccccaccatggcttttatagctgaagatgatg (IVTTIL18-25), gaaattaa-tacgactcactatagggagaccccaccatggctgatgatgaaaacctggaat (IVTTIL18-29), gaaattaatacgactcactataggga-gaccccaccatggctctggaatcagattactttg (IVTTIL18-33), gaaattaatacgactcactatagggagaccccaccatggcttactttgg-caagcttgaat (IVTTIL18-37), gaaattaatacgactcactatagggagaccccaccatggctcttgaatctaaattatcag (IVTTIL18-41), gaaattaatacgactcactatagggagaccccaccatggctataagaaatttgaatgacc (IVTTIL18-48), gaaattaatacgactcac-tatagggagaccccaccatggctattgaccaaggaaatcggc (IVTTIL18-58), in combination with 3’ primer ttttttttttttttttttctagtcttcgttttgaacag (IVTTIL18-term). Loading for IL-18R reporter assays was normalized by total IL-18 measured by ELISA (R&D Systems) following manufacturer protocol.

### Ethics Statement

This study was conducted according to the principles expressed in the Declaration of Helsinki. Blood was collected from healthy adult volunteers under informed consent and approved by the Institutional Review Board of Emory University. Animal experiments were approved by the Institutional Animal Care and Use Committees of Emory University.

### Statistics and Data Analysis

Values are expressed as mean ± standard deviation. Differences between groups were analyzed using 1-way analysis of variance with Dunnett multiple comparisons analysis unless otherwise indicated. Differences are considered statistically significant at a P value of <.05 using GraphPad Prism v9 software. Principal Component Analysis (PCA) was used to reduce the number of variables needed to adequately describe differences in cytokine profiles; each condition was treated as an independent variable and multivariate analysis was performed in Prism v9. Protein models were visualized using PyMol 2.3.3. Diagrams created with BioRender.com.

## Acknowledgements

We thank Victor Nizet for his insights and bacteria, all members of the LaRock lab for discussions, Cassandra Quave for cell lines, the Children’s Healthcare of Atlanta and Emory University’s Children’s Clinical and Translational Discovery Core (CTDC) for whole blood and processing, Shonna McBride, BEI Resources, and the Emory Investigational Clinical Microbiology Core (ICMC) for bacterial isolates.

## Author contributions

C.N.L. conceived this study, C.N.L., A.F.J., and J.S.S. designed the experiments and wrote the paper. A.F.J., J.S.S., K.T., R.R., D.L.L., and C.N.L. performed experiments and analyzed results.

